# Comparative spatial transcriptomics of peach and nectarine fruits elucidates the mechanism underlying fruit trichome development

**DOI:** 10.1101/2023.10.03.560746

**Authors:** Zihao Zhao, Ke Cao, Aizhi Qin, Zhixin Liu, Liping Guan, Susu Sun, Hao Liu, Yaping Zhou, Jincheng Yang, Yumeng Liu, Mengke Hu, Vincent Ninkuu, Xuwu Sun, Lirong Wang

## Abstract

Peach (*Prunus persica*) is an economically significant fruit-bearing tree. The trichomes of peach fruits are essential for their growth and development by playing a protective role against abiotic stresses, such as intense ultraviolet (UV) light, high temperature, cold-induced stress, and biotic stresses, such as pests and disease infestation. However, the mechanism underlying trichome development in peach fruits is unknown. Therefore, spatial transcriptome sequencing technology was used to compare the transcriptomic information seven days after flowering (DAF) of peach and nectarine fruits at an ultra-high resolution. The results revealed significant variations in how fruits at the early development stage responded to stress exposure. Comparatively, nectarine response to stress was significantly magnified than peach. Notably, a novel trichome-related marker gene, *Prupe.7G196500*, was identified in peach, which showed a robust function in response to jasmonic acid and wounding. Further, the *Prupe.7G196500pro::GUS* construct was found to specifically expressed in the trichomes of the true leaves of 10-day-old *Arabidopsis thaliana* (*Arabidopsis*) seedlings and induced by drought treatment. Finally, gain-of-function analysis showed that *Prupe.7G196500* promoted trichomes development. In conclusion, this study identified the cell types and novel marker genes associated with peach fruit. The role of *Prupe.7G196500* in positively regulating trichome development was also characterized, thereby laying the foundation for analyzing the mechanism involved in trichome development in peach fruits.

## INTRODUCTION

Peach (*Prunus persica*) is an economically valuable fruit and an essential source of phenolics, cyanogenic glucosides, and phytoestrogens, which intimately affect human health (Rodriguez et al., 2019). The peach fruit has a hairy outermost exocarp protecting the edible fleshy mesocarp, which harbors the hard endocarp (Dardick and Callahan, 2014). Peach is classified as a drupe because its endocarp is lignified during development (Guo et al., 2018). The fuzzy skin on peach fruits is a characteristic of pubescence, known as fruit trichomes (Fernandez et al., 2011). Trichomes are highly differentiated from epidermal cells that are widely distributed on the surface of various organs/tissues and play a critical role in the evolution of plants (Szymanski et al., 2000). Depending on their characteristics and functions, trichomes are unicellular or multicellular, branched or unbranched, and glandular or non-glandular (Arteaga et al., 2021; Wang et al., 2021). In addition, trichomes occur in a variety of shapes, such as head, star, hook, scale, etc., (Arteaga *et al*., 2021; Wang *et al*., 2021). The trichomes of peach fruits are unicellular, non-glandular, and play a protective role during the stages of early development and also ripening. They first appear on the surface of the ovary four weeks before flowering, and mostly dies when the fruit matures physiologically (Fernandez *et al*., 2011).

Plants have evolved various defense mechanisms against the biotic and abiotic stresses occurring in the environment that they may be exposed to (Hauser et al., 2001). Trichomes are the first defense barriers to pathogen invasions and feeding herbivore attacks (Santamaria et al., 2013). Plants interact with the environment by regulating the changes in the shape and density of their trichomes, thus affecting their physiological processes (Wagner et al., 2004). Glandular trichomes with secretory functions are organs that are involved in the synthesis of biochemicals and secretion of a variety of defense-related compounds (Schuurink and Tissier, 2020), such as terpenoids (Bruckner et al., 2014; Sallaud et al., 2009; Schilmiller et al., 2009), flavonoids (Kim et al., 2014; Schmidt et al., 2011; Tattini et al., 2000), acylsugars (Fan et al., 2016; Schilmiller et al., 2012; Schilmiller et al., 2015), etc. For instance, the trichomes of tea plants (*Camellia sinensis* L.) can synthesize an array of defense metabolites, including catechins, theanine, caffeine, flavonols, saponins, terpenes, and lipid volatiles, which demonstrate either toxicity or inhibitory effects against herbivores and pathogens (Li et al., 2020). In addition, they are also highly beneficial for the development and growth of fruits by attracting beneficial insects and microorganisms but inhibit the germination and growth of competing plants (Howe and Jander, 2008; Leckie et al., 2016; Massalha et al., 2017). Non-glandular trichomes, such as the densely soft trichomes of young peach fruits or the bramble or prickly trichomes of cucumber (*Cucumis sativus* L.) (Du et al., 2020; Liu et al., 2018; Zhang et al., 2016), are also crucially involved in physical defenses by using their morphological structures to distract herbivores (Gangasaran et al., 2010).

Trichomes also play a significant role in mitigating abiotic stresses in plants (Zhao and Chen, 2016). The presence of trichomes increased the thickness of the epidermal cells and markedly enhanced the long-chain fatty acid contents compared to those of the other epidermal cells, which play a significant role in regulating the temperature and reducing transpiration (Busta et al., 2017; Hegebarth et al., 2016). In alpine-rich areas, *Croton tiglium* and *Vriesea* absorb atmospheric water and nutrients through their trichomes, significantly improving the water- and fertilizer-use efficiencies (Vanhoutte et al., 2017; Vitarelli et al., 2016). The dense, multi-branched, spine-like trichomes provide strong resistance against the sand blown along with the winds and can help reduce the mechanical damage caused by them (Chen et al., 2014). Furthermore, the trichomes of the aquatic plant *Salvinia molesta* play a hydrophobic role in maintaining normal respiration (Barthlott et al., 2009). The glandular trichomes play an essential role in helping plants adapt to low temperature-induced stress through flavonols-(Bhatia et al., 2018) and terpenoids-mediated mechanisms (Koudounas et al., 2015). In addition, polyamines secreted by trichomes interact with abscisic acid (ABA) metabolic pathways to activate the production of reactive oxygen species (ROS) and nitric oxide (NO), which regulate the state of the ion channels and Ca^2+^ homeostasis, thus conferring a protective role in response to various abiotic stresses (Diao et al., 2017; Pottosin et al., 2014). Similarly, mono- and sesquiterpenoids secreted by the glandular trichomes of *Artemisia annua L.* increase after drought-induced stress, suggesting that volatile terpenoids may play a role in regulating drought resistance (Yadav et al., 2014). Additionally, trichomes can also physically resist abiotic stresses. Densely spaced trichomes can effectively mitigate the effects of direct sunlight rays, thus preventing water loss (Koudounas *et al*., 2015), and mitigate the effect of high temperature, UV radiation, and drought-induced stress (Karabourniotis et al., 2020). These mechanisms indirectly affect water-use efficiency and photosynthetic and transpiration efficiencies of plants (Bickford, 2016). Trichomes also help maintain ion homeostasis; notably, the Solanaceae family members are crucially involved in detoxifying heavy metals (Cd, Ni, Pb, and Zn) (Koul et al., 2021).

In recent years, significant progress has been made in understanding the molecular mechanisms behind trichome formation and development in peach fruits. Earlier studies suggested that the peach/nectarine character was monogenic (G/g), and the furless traits of nectarines were recessive to the furry traits of peaches (Vendramin et al., 2014). The “G” locus was mapped to the distal part of the linkage group (LG) 5 (Dirlewanger et al., 2007; Le Dantec et al., 2010) spanning a region, 1.189 Mb long, from 15,126,681 to 16,315,341 of the pseudomolecule 5 of the peach reference genome (Peach v1.0) (Verde et al., 2013). The gene encoding a member of the MYB family of transcription factors (TFs), *PpMYB25,* positively regulates trichome formation in peach fruits (Vendramin *et al*., 2014). The insertion of the Ty1-copia retrotransposon within the third exon of *PpMYB25* was identified as the putative cause of the glabrous phenotype of nectarine fruits (Vendramin *et al*., 2014). *PpMYB26*, a homolog and a downstream protein of *PpMYB25*, is also involved in regulating the formation of trichomes in fruits (Yang et al., 2022). It is well known that multiple genes regulate several critical developmental processes in plants. For example, the formation of the unicellular, non-glandular trichomes in *Arabidopsis* is controlled by the R2R3-MYB/bHLH/WD40 repeat complex, which activates the transcription of the HD-ZIP-encoding *GLABRA2* (*GL2*) gene (Oppenheimer et al., 1991; Zhao et al., 2008). Similarly, the formation of trichomes in peaches may also be regulated by several unknown transcriptional networks, which need further investigation.

The emerging technology of spatial transcriptomics (ST) allows for studying the expression profiles and spatial distribution of genes *in situ* at an ultra-high resolution (Rao et al., 2021). ST can generate the entire transcriptomic data from a complete tissue sample and preserve the spatial background information regarding the gene expression patterns while resolving the gene expression profiles (Stahl et al., 2016). The application of this technology in plants mainly focuses on constructing spatiotemporal maps, defining new types of tissues/organs, identifying and discovering novel putative marker genes, studying the dynamics of tissue and organ development, and analyzing gene regulatory networks (Giacomello, 2021).

Although limited progress has been made in understanding the molecular mechanism underlying trichome formation in peaches, it is still largely undetermined. In this study, ST sequencing of peach and nectarine fruits seven days after flowering (DAF) was conducted, and the cell types of the two were compared. Finally, the differentially expressed genes (DEGs) in each cell type were screened and identified. Further, the transcriptional regulatory networks of the DEGs identified in nectarine and peach fruits were ascertained. Based on the results, a marker gene for peach trichomes, *Prupe.7G196500,* was identified. The ectopic expression of *Prupe.7G196500* in *Arabidopsis* showed that it was expressed in the trichomes, regulated trichome development, and modulated environmental stress responses. The successful application of ST in peach fruits provides a unique opportunity to analyze the formation and development of trichomes.

## RESULTS

### Spatial gene expression profiles of nectarine and peach fruit cell populations

The distinguishing feature between nectarine and peach fruits is the presence of a smooth glabrous exocarp in nectarine and densely spaced trichomes in peach. Previous studies have shown that plant trichomes are important in providing resistance against various stresses (Bhatia *et al*., 2018; Santamaria *et al*., 2013). However, the regulatory network behind trichome formation in peaches is still unclear. Therefore, the spatial gene expression profiles of cell populations of nectarine and peach fruits at seven DAF were ascertained and compared to further analyze the regulatory network underlying trichome formation (Figure 1).

**Figure 1.**
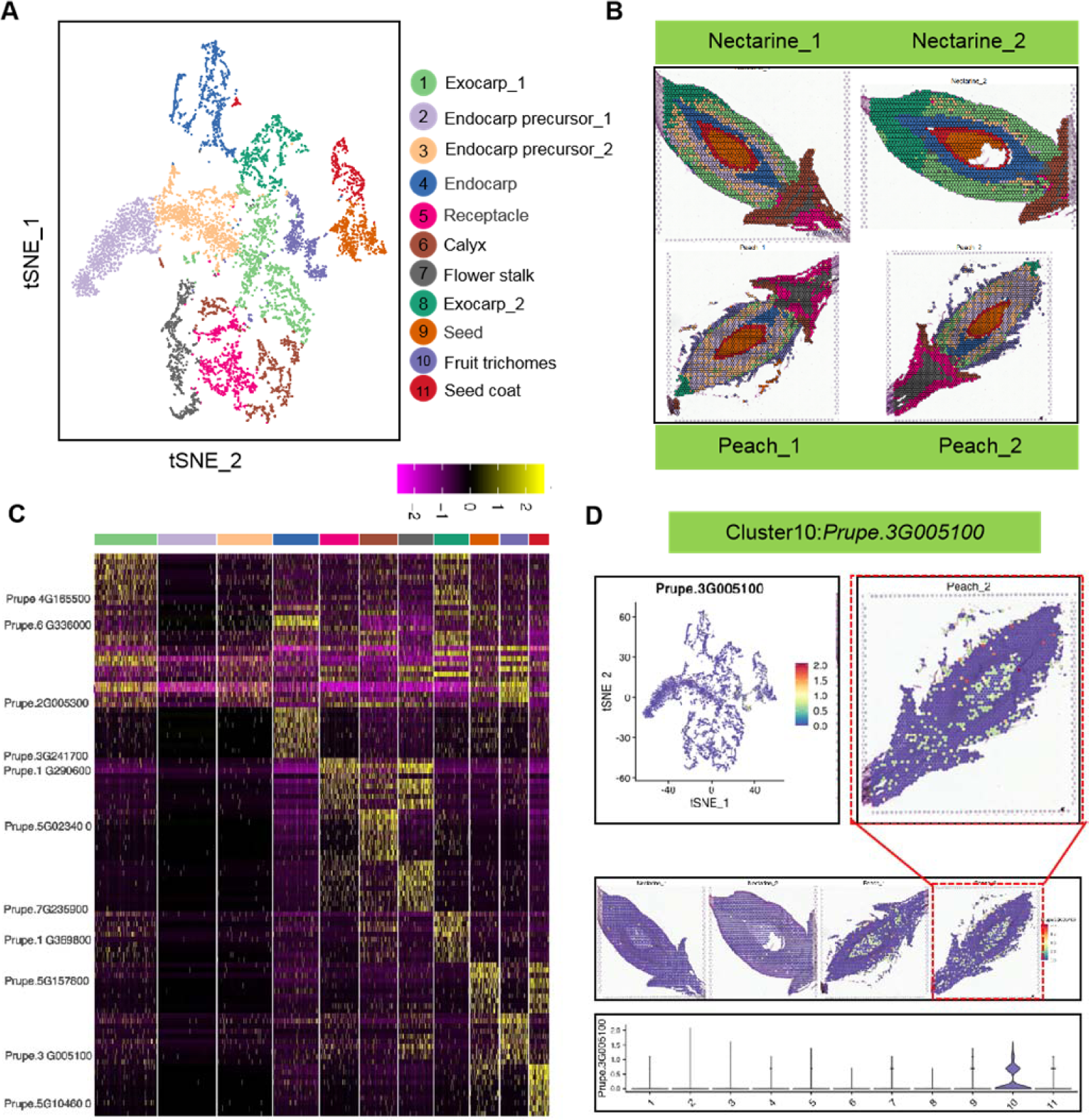
Spatial transcriptomics analysis of Nectarine_1, Nectarine_2, Peach_1, and Peach_2. **(A)** t-SNE analysis of Nectarine_1, Nectarine_2, Peach_1, and Peach_2. The number in each figure represents a different cell type, and various colors represent the different cell types. **(B)** All clusters in **(A)** were mapped to their spatial locations. The results indicated that the 11 clusters were located in different organizational regions. **(C)** Heatmap of the expression patterns of the marker genes representative of each cell cluster. The ordinate represents the marker genes in each cell cluster, the colors varying from purple to yellow represent the expression levels of the marker genes from low to high, and the abscissa represents the different cell clusters. **(D)** The spatial expression patterns of the representative marker gene of cluster 10. The related to the t-SNE and gene is expressed as the violin map.

Spatial transcriptomics (ST) sequencing was performed on one nectarine cultivar: Nectarine_1 and Nectarine_2, and one peach cultivar: Peach_1 and Peach_2, at seven DAF. After further screening the sequencing data for acceptable levels of quality, the total number of genes identified in the Nectarine_1, Nectarine_2, Peach_1, and Peach_2 samples were 19385, 19654, 18817, and 18894, respectively. The high-quality spots obtained per sample section were 2000, 1873, 1906, and 1979, respectively. Specifically, the average number of genes in each spot was 3320, 4756, 2498, and 2739; the average number of unique molecular identifiers (UMIs) in each spot was 10756, 16963, 8571, and 9332; and the average proportion of mitochondrial genes in each spot was close to 0% (Supplemental Figure S1A – C, Supplemental Table S1).

The t-distributed Stochastic Neighbor Embedding (t-SNE) algorithm (van der Maaten, 2014) was used to apportion all spots into 11 clusters, each labeled with a different color, based on the gene expression similarities observed between the four sample sections (Figure 1A). Subsequently, the spatial barcode information carried on the spots was used to restore these spots to the original location of the sample tissue section (Figure 1B). The clustering results revealed that the composition of the cell populations of nectarine and peach fruits was similar, except for cluster 10, which was unique to peach and appeared only in the Peach_1 and Peach_2 samples (Figure 1B, Supplemental Figure S1E). The cell types’ annotated results for the 11 cell clusters included: cluster 1 was associated with exocarp_1, cluster 2 with endocarp precursor_1, cluster 3 with endocarp precursor_2, cluster 4 with endocarp, cluster 5 with receptacle, cluster 6 with calyx, cluster 7 with flower stalk, cluster 8 with exocarp_2, cluster 9 with seed, cluster 10 with fruit trichomes, and cluster 11 with seed coat. Interestingly, the samples of peach_1, peach_2, nectarine_1, and nectarine_2 were observed in turn, and it was found that the endocarp (cluster 4) developed and formed gradually, with a tendency to progressively wrap the seed and then the seed coat (Figure 1B).

To characterize the novel marker genes associated with the identified cell populations, a heatmap indicating the expression levels of the ten topmost genes enriched in each cluster was conducted (Figure 1C). The spatial expression profiles of the potential marker genes with high expression in each cluster were ascertained, including some with known functions (Figure 1D and Supplemental Figure S2). For example, *Prupe.2G005300*, a marker gene for endocarp precursor_2 (cluster 3), and its homolog in *Arabidopsis*, *AtLOX2,* encodes an isoenzyme of lipoxygenase, which played a vital role in the biosynthesis of the plant growth regulators (JA and ABA) (Bell et al., 1995). The marker gene for endocarp (cluster 4) was identified as *Prupe.3G241700*, which was determined to be involved in lignin biosynthesis (Supplemental Table S2). It has been proven that, during the development of peach fruit, the endocarp undergoes gradual lignification to form pits (Guo *et al*., 2018). This study suggested that *Prupe.2G240500* encoded a mannose-1-phosphate guanylyltransferase, expressed in clusters 3 and 4 (Supplemental Table S2). *Prupe.6G116900* encoded a UDP-glucuronate decarboxylase, expressed in clusters 2 and 4 (Supplemental Table S2). *Prupe.2G240500* (*ppa007618m*) and *Prupe.6G116900* (*ppa006917m*) were expressed in the endocarps of peach fruits, nine DAF (Rodriguez *et al*., 2019). *Prupe.1G290600* was recognized as a marker gene of the receptacle (cluster 5); its homolog in *Arabidopsis*, *AtAPETALA1* (*AtAP1*), plays a crucial role in flower initiation and regulate the expression of B-class genes that control stamen development (Ondar et al., 2008). *Prupe.5G023400* was identified as a marker gene for the calyx (cluster 6); its homolog in *Arabidopsis*, *AtPRR1,* played a role in flowering and early photomorphogenesis (Matsushika et al., 2007). *Prupe.7G235900* was ascertained to be a marker gene for flower stalk (cluster 7); its homolog in *Arabidopsis*, *AtEFM,* mediated flowering response to environmental cues (Yan et al., 2014). *Prupe.1G369800* was determined to be a marker gene of exocarp_2 (cluster 8); its homolog in *Arabidopsis*, *AtGOXL3,* encoded a galactose oxidase, which crosslinks pectin molecules by modifying the galactose side chain and promoted intercellular adhesion between epidermis cells (Sola et al., 2021). *Prupe.5G157800* was also recognized as a marker gene of seeds (cluster 9), and its homolog in *Arabidopsis*, *AtSFAR4,* encoded a GDSL-type esterase, which enhanced the expression of genes related to fatty acid metabolism during seed germination (Huang et al., 2015). In addition, *Prupe.3G005100* was identified as a marker gene for fruit trichomes (cluster 10), and its cotton homolog, *GhGolS1*, encoded a galactinol synthase that is involved in the raffinose biosynthesis, an essential metabolite that improves the quality of cotton fibers (Zhou et al., 2012). The marker gene associated with seed coat (cluster 11) was identified as *Prupe.5G104600* and its homolog in *Arabidopsis*, *AtBEL1,* promoted ovule development (Western and Haughn, 1999).

### Validation of the expression patterns of the marker genes by *in situ* hybridization (ISH)

To verify the reliability of the definitions of the cell cluster made based on the ST data, RNA ISH was performed on the novel exocarp (*Prupe.6G012100*) and endocarp (*Prupe.8G228000*) marker genes. The results suggested that *Prupe.6G012100* was expressed in the exocarp and *Prupe.8G228000* in the endocarp of the nectarine and peach fruits in consistencewith those observed by the ST (Figure 2, Supplemental Figure S3). These results confirmed the reliability of the definitions of the cell clusters and the availability of the new marker genes made in this study.

**Figure 2.**
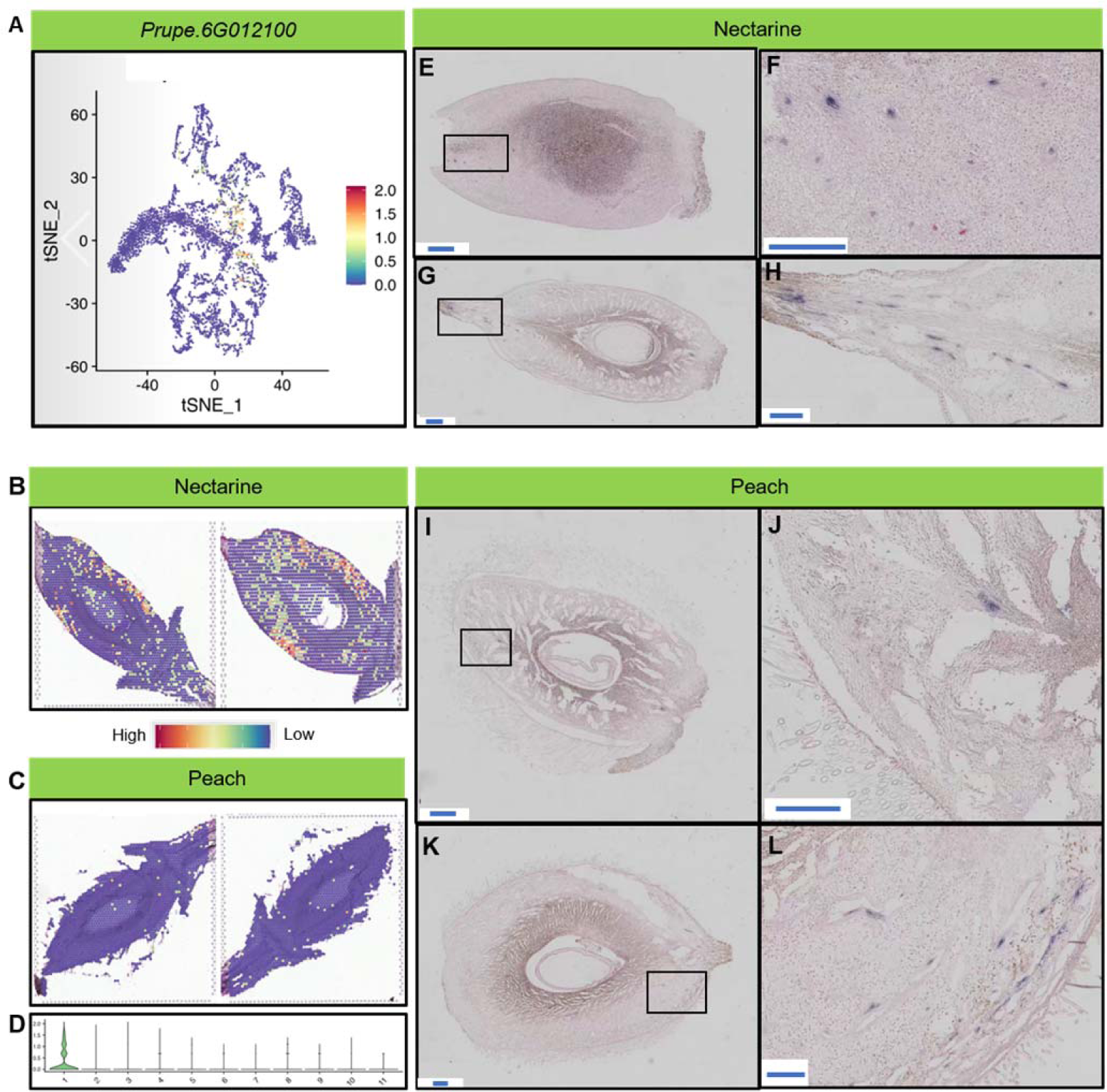
Validation of gene expression by ISH. **(A – D)** Spatial location map and the violin plot indicating the expression levels of the exocarp marker gene, *Prupe.6G012100,* in the different clusters of the ST data. **(E – H)** Tissue sections from nectarine fruits analyzed by ISH illustrate the spatial distribution of *Prupe.6G012100*. **(I – L)** Tissue sections from peach fruits analyzed by ISH, illustrate the spatial distribution of *Prupe.6G012100*. Scale bar: 500 µm for E, G, I, and K; 250 µm for F, H, J, and L.

### Screening of the marker genes associated with fruit trichomes

To enrich the available marker genes resources associated with peach trichomes, the spatial expression characteristics of the ten topmost highly expressed genes in the trichomes (cluster10) were characterized (Figure 3). The results revealed that these genes were highly and specifically expressed during the early development of trichomes, suggesting they could be used as markers for screening and identifying trichomes in peach fruits. These genes are reported to participate in abiotic stress responses in other plant species. For example, the *Arabidopsis* gene *AtELIP1*, a homolog of *Prupe.1G021800*, protects photosynthetic components against photooxidation-induced stress (Rossini et al., 2006). In addition, the *Prupe.2G218700* homolog in *Arabidopsis* (*AtRCI2C*) mediates salt-induced stress (Kim et al., 2016; Long et al., 2015), while the homolog of *Prupe.3G005100*, *AtGolS1*, confer resistance against heat-induced stress (Panikulangara et al., 2004). Furthermore, *Prupe.3G079700* could be functionally similar to its *Arabidopsis* homolog, *AtNAD-ME1*, which is involved in the oxidation of L-malate in the mitochondria (Tronconi et al., 2010). Similarly, the *Arabidopsis AtHB13*, a homolog of *Prupe.4G041900*, conferred tolerance to cold stress by inducing the accumulation of pathogenesis-related proteins and glucanases (Cabello et al., 2012). The *Arabidopsis AtA6PR1* is also a homolog of *Prupe.8G083400*, is reported to be responsive to abiotic stresses (Rojas et al., 2019). The *Arabidopsis* gene *AtRAB28*, a homolog of *Prupe.8G238700*, could be induced by ABA (Busk et al., 1999). However, the potential functions of these marker genes in regulating the formation of trichomes in peach fruits need further elucidation.

**Figure 3.**
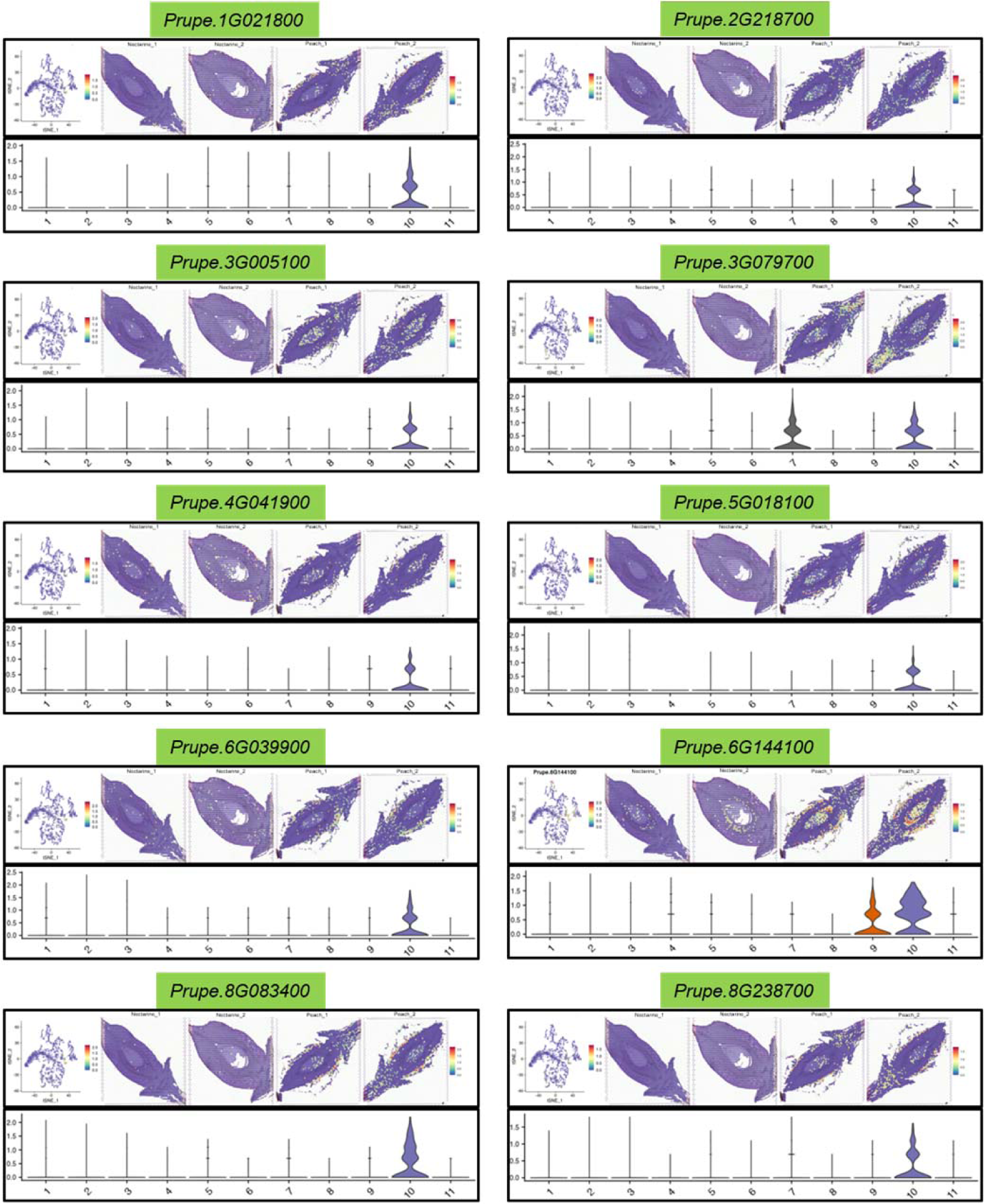
Representative marker genes identified in the fruit trichomes. The figure shows ten marker genes that were highly expressed and with tissue-specificity in cluster 10. The expression of these genes ranges from low to high, as indicated by blue to red in the spatial visualization. The violin diagram shows the expression site of the gene in the tissues.

### Gene Ontology (GO) enrichment analysis of nectarine and peach fruits

Nectarine fruit possesses smooth and glossy skin, while peach fruit is covered by pubescence, an apparent phenotypic difference linked to the variations in the expression of trichome formation-related genes during fruit development. To identify the genes crucial for trichome formation, DEGs identified in the 11 clusters based on the ST data from nectarine and peach fruits were screened (Supplemental Table S2). Through GO enrichment analysis of the DEGs in the different clusters, the metabolic processes and functions associated with these clusters were determined. Endocarp precursor_1 (cluster 2), endocarp precursor_2 (cluster 3), and endocarp (cluster 4) were all highly enriched under similar GO terms, indicating the reliability of the cell cluster annotations made in this study. These three clusters are involved in “carbohydrate-mediated signaling pathways” and “cellular polysaccharides metabolic processes” (Figure 1E).

To analyze whether nectarine and peach fruits differed only in the early stage of development, GO enrichment analysis and comparison of the DEGs identified between the two was conducted. Compared with peach, the GO term enriched by the up-regulated DEGs in nectarine fruits were mainly related to biotic stimulus and defense responses (Figure 4A), suggesting the loss of trichomes of nectarine fruits could compromise stress responsiveness. In addition, the up-regulated DEGs in nectarine were highly enriched under “saponin biosynthesis process,” “flavonoid biosynthesis process,” and “sterol metabolic process” (Figure 4A). This may be due to the ability of nectarine to resist environmental stresses after the loss of trichomes by increasing the accumulation of defense-associated metabolites such as saponins (Thimmappa et al., 2014), flavonoids (Kim *et al*., 2014; Schmidt *et al*., 2011; Tattini *et al*., 2000), and sterols (Clouse, 2002). Compared with peach fruits, the GO terms enriched by down-regulated DEGs in nectarine fruits included “response to water deprivation,” “abscisic acid-activated signaling pathway,” and “response to abscisic acid” (Figure 4A). These results indicated that nectarine and peach fruit were markedly different in the early stage of development, and the expression levels of genes related to environmental stress tolerance in nectarine were significantly enhanced compared with peach fruit. This may be due to the loss of trichomes of nectarine fruits during evolution, suggesting nectarine could be more susceptible to stresses than peach fruits. In addition, the expression levels of drought-stress-related genes in nectarine fruits were significantly reduced thanpeach fruits, suggesting that nectarine had a more robust water retention capacity than peach. The possible reason for this phenomenon may be the thick cuticle covering the exocarp in nectarine fruits (Kosma et al., 2009; Shepherd and Griffiths, 2006).

**Figure 4.**
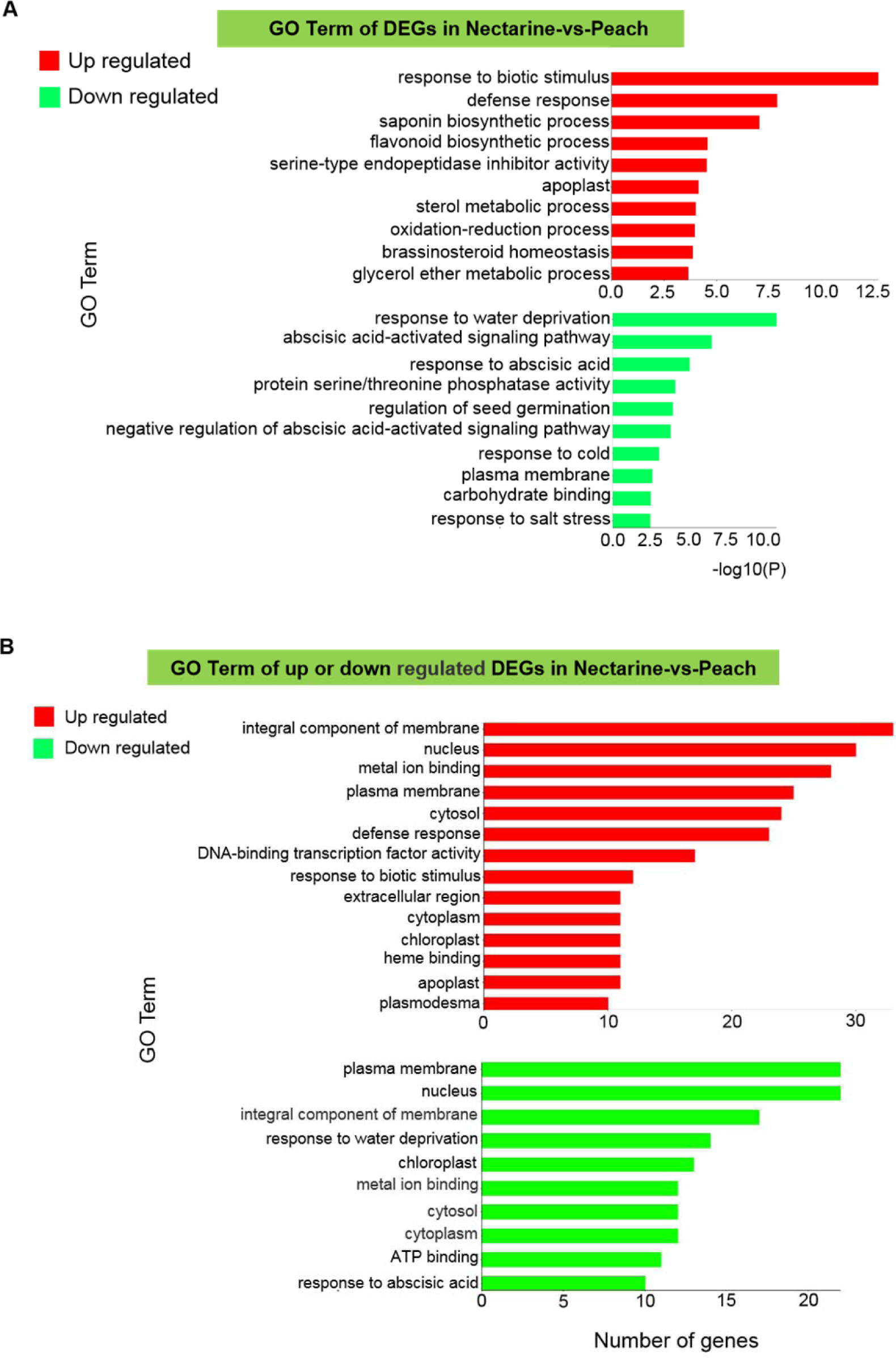
GO enrichment analysis in nectarine relative to peach. (A) GO enrichment map of up- or down-regulated differentially expressed genes (DEGs) in nectarine relative to peach. (B) The number of the significantly up- or down-regulated DEGs in nectarine relative to peach.

Furthermore, the number of genes enriched under the GO terms was used to ascribe the reason underlying nectarine being more liable to stresses than peaches under adverse conditions. The results revealed that the number of up- and down-regulated genes in nectarine were higher than in peach and enriched “response to stimulus,” “metabolic process,” and “catalytic activity” GO terms (Supplemental Figure S4A). This phenomenon in nectarine may be due to the loss of trichomes during evolution, enhancing its susceptibility to stresses. Evidence of conspicuous changes in the metabolic and catalytic processes associated with this phenomenon in plants under abiotic stress are reported (Cardoso et al.; Khan et al., 2019).

Further, 216 significantly up-regulated and 160 remarkably down-regulated DEGs were identified in nectarine compared with peach (Supplemental Table S3). GO enrichment analysis on these DEGs was then performed, and the results only preserved the GO terms associated with more than ten genes (Figure 4B, Supplemental Table S3). The proportion of up-regulated DEGs involved in “defense response” and “response to biotic stimulus” was higher. Conversely, that of down-regulated DEGs involved in “response to water deprivation” and “response to abscisic acid” was higher (Figure 4B). In addition, the heatmap analysis of the DEGs-enriched GO terms demonstrated that the DEGs involved in “response to water deprivation” were down-regulated. In contrast, those mediating “defense response to other organisms,” “response to wounding,” and “flavonoid biosynthetic process” were up-regulated in nectarine compared with peach (Supplemental Figure S4B). Following this, the biological process and molecular function enrichment analysis for these DEGs were performed, and the directed acyclic graphs for the enriched terms were plotted. Compared with peach fruits, the up-regulated DEGs in nectarine fruits were highly enriched in “response to stress,” “defense response,” “steroid metabolic process,” and “saponin metabolic process” (Supplemental Figure S5), suggesting that defense- and metabolism-related processes were more predominant in nectarine than in peach fruits. Moreover, the up-regulated DEGs in nectarine fruits were highly enriched in “oxidoreductase activity” and “anion transmembrane transporter activity” than those in peach fruits (Supplemental Figure S6). Previous studies have shown that anions are essential in energy metabolism and metabolite transport (Li et al., 2013). Therefore, it can be concluded that nectarine is more sensitive to environmental stress but more resistant to water deprivation than peach.

In summary, a comparison of multiple parameters between nectarine and peach fruits suggested that after the loss of trichomes of nectarine fruits, the exocarp was directly exposed to the environmental conditions, resulting in higher activity of defense-related metabolic pathways and up-regulation of genes related to stress-resistance. This indirectly proved that trichomes participate in stress resistance Perhaps this phenomenon could be due to the thick protective cuticle covering the exocarp of the nectarine fruits, shielding it from water loss (Kosma *et al*., 2009; Shepherd and Griffiths, 2006).

### Analysis of the transcription factors (TFs) regulatory networks associated with identified DEGs in nectarine and peach

A species-specificity in the TF-regulatory networks may indicate the occurrence of different developmental processes between species. Therefore, the accuracy of the results of our analysis was verified by exploring the differences in the TFs detected in nectarine and peach during fruit development. A regulatory network analysis of TFs associated with all the DEGs identified between nectarine and peach was performed, resulting in the identification of nine key TFs, including three WRKY, three AP2/ERF-ERF, two bZIP, and one HB-HD-ZIP TFs (Figure 5, Supplemental Table S4). All four TFs have been reported to be essential in responding to biotic and abiotic stresses (Ambawat et al., 2013; Droge-Laser et al., 2018; Erpen et al., 2018; Wani et al., 2021; Xie et al., 2019).

**Figure 5.**
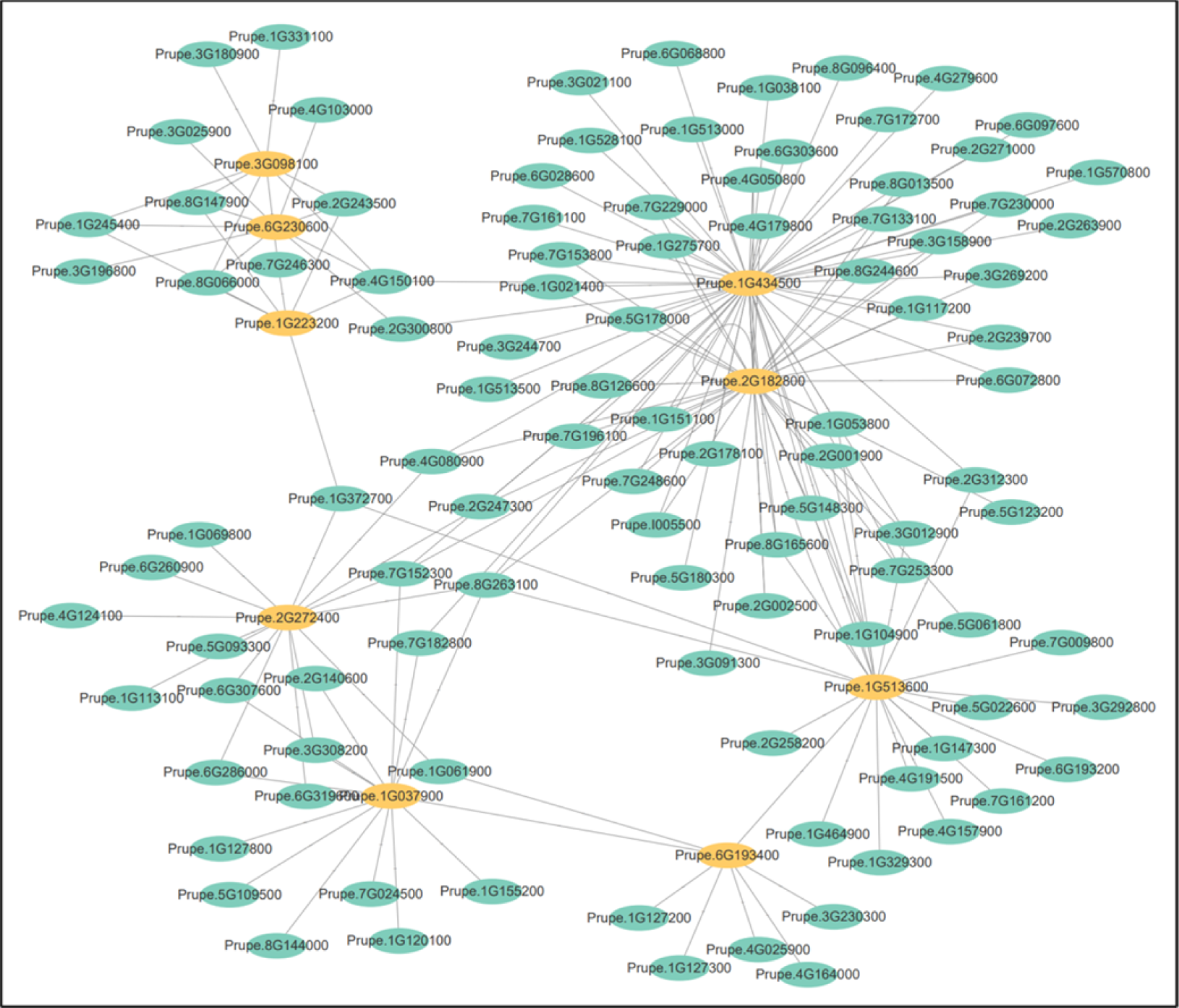
The map of the transcription factor (TF) regulatory network for the differentially expressed genes in nectarine and peach. The TFs with the yellow background are crucial to the regulatory network. These included three WRKY TFs (Prupe.3G098100, Prupe.6G230600, and Prupe.1G223200), three AP2/ERF-ERF TFs (Prupe.2G272400, Prupe.1G037900, and Prupe.1G513600), two bZIP TFs (Prupe.1G434500 and Prupe.2G182800), and one HB-HD-ZIP TFs (Prupe.6G193400).

Specifically, the three WRKY TFs identified were *Prupe.3G098100* (*PpWRKY40*), *Prupe.6G230600* (*PpWRKY7*), and *Prupe.1G223200* (*PpWRKY75*). The *Arabidopsis AtWRKY40* is a homolog of *Prupe.3G098100*, which complementsWRKY18 and 60 to coordinate responses to ABA and abiotic stress (Chen et al., 2010). In addition, the *Arabidopsis* TFs WRKY40 and 18 negatively regulate flg22-induced genes, thereby preventing exaggerated defense responses (Birkenbihl et al., 2017). The *Arabidopsis AtWRKY7*, a homolog of *Prupe.6G230600* plays a negative role in the defense response against *Pseudomonas syringae* (Kim et al., 2006), while *AtWRKY75* in *Arabidopsis,* which is also a homolog of *Prupe.1G223200*mediates jasmonate (JA) synthesis against necrotrophic fungal pathogens in plants (Chen et al., 2021). The three AP2/ERF-ERF TFs identified were *Prupe.2G272400* (*PpERF105*), *Prupe.1G037900* (*PpERF1*), and *Prupe.1G513600* (*PpRAP2.4*). The *Prupe.2G272400* is a homolog of *Arabidopsis AtERF105*, which regulates cold-induced stress (Illgen et al., 2020) and defense against *P. syringae* (Cao et al., 2019). Further, the *Arabidopsis* ETHYLENE RESPONSE FACTOR1 (ERF1), an upstream component of JA and ethylene (ET) signaling pathways, and participates in pathogen resistance and response to salt-induced stress (Cheng et al., 2013; Huang et al., 2016) is homologous to *Prupe.1G037900*. Also, the *Arabidopsis AtRAP2.4*, which is homologous to *Prupe.1G513600* is responsive to salt-and drought-induced stresses (Lin et al., 2008). In addition, the *AtRAP2.4* TF activates cuticular wax biosynthesis of in *Arabidopsis* leaves under drought-induced stress (Yang et al., 2020).

The two bZIP TFs identified included *Prupe.1G434500* (*PpABF2*) and *Prupe.2G182800* (*PpGBF3*). The *Arabidopsis AtABF2* was identified as a homolog of *Prupe.1G434500* predominantly acts downstream of SRK2D/E/I in the ABA signaling pathway in response to osmotic stress during vegetative growth (Yoshida et al., 2015). ABF2 TF also interacts with the NAC TF, ANAC096, in response to dehydration- and osmotic-induced stress (Xu et al., 2013). The *AtGBF3*, which crucially induces drought tolerance in *Arabidopsis* (Ramegowda et al., 2017) is homologous to *Prupe.2G182800*. Similarly, the *Arabidopsis* AtHB6 is homologous to the HB-HD-ZIP TF encoded by *Prupe.6G193400*, which may be involved in the ABA signaling pathways (Liu et al., 2011). These results indicated that nectarine and peach possess different developmental processes and that WRKY, AP2/ERF-ERF, bZIP, and HB-HD-ZIP TFs play a crucial role in stress response.

### *Prupe.7G196500* positively regulates the development of trichomes

Based on the GO enrichment analysis of the DEGs recognized in fruit trichomes (cluster 10), *Prupe.7G196500* was identified as a putative gene with a potential function in regulating trichome development. We found that *Prupe.7G196500* was enriched in the GO terms mainly for “response to wounding” and “response to jasmonic acid” (Supplemental Table S2). Whereas exogenous application of JA promoted trichome formation on the leaf surfaces of Rhodes Grass (*Chloris gayana* Kunth) (Kobayashi et al., 2010), in *Arabidopsis*, JAs treatment induced the degradation of jasmonate-ZIM-domain (JAZ) proteins to activate the WD-repeat/bHLH/MYB complex for trichome formation (Qi et al., 2011).

The search for *Prupe.7G196500* homolog in *Arabidopsis* identified *AtSSL4* (*AT3G51420.1*) and *AtSSL5* (*AT3G51430*) with the highest homology that encodes: strictosidine synthase-like (SSL) proteins and plant defense signaling compounds such as salicylic acid and methyl jasmonate induced the expression of *AtSSL5*, but not *AtSSL4*, indicating that *AtSSL4* played a specific role in the innate, while *AtSSL5* in the inducible defense responses in plants (Sohani et al., 2009). Further, phylogenetic analysis of *Prupe.7G196500* suggested that it was closely related to *AtSSL5* (Supplemental Figure S7). Therefore, it can be speculated that *Prupe.7G196500* may possess functions similar to *AtSSL4* and *5* for trichomes development in peach fruits by participating in the JA-responsive pathway, thereby inducing resistance in immature fruits against biotic and abiotic stresses.

To determine the function of *Prupe.7G196500* in peach, the differences in its expression levels in peach and nectarine were analyzed. Based on the previous pan-genomic data, the genotypes of 565 peach and 179 nectarine germplasms were identified. Seven loci determined to be mutations were mapped in the mRNA and promoter of *Prupe.7G196500*. A correlation analysis of these mutations and the corresponding phenotypes showed that the SNP mapped to the 3’-UTR (Chr7:18,561,307 bp) most correlated with trichome development in peach, with a p-value of 0.05 (Figure 6A). Using peach germplasm “Kashi No. 1” as a reference material, the expression patterns of *Prupe.7G196500* in different tissues of the peach plant tissues were analyzed. The results revealed that *Prupe.7G196500* was mainly expressed in the fruits (Figure 6B). The expression patterns of *Prupe.7G196500* ascertained in different tissues of peach and nectarine fruits at seven DAF demonstrated that *Prupe.7G196500* was highly expressed in the trichomes, partially in the peel mixture of peach fruit but almost none-responsive in the peel mixture of nectarine fruits (Figure 6C). Next, we analyzed the expression patterns of *Prupe.7G196500* in the peach germplasm “Zhengbai 5-2” at different fruit development stages, which revealed that *Prupe.7G196500* transcribes mainly accumulated in the early fruit development stage (Figure 6D). The possible correlation between trichome length and *Prupe.7G196500* expression levels during the ripening process of hairy peach fruits was explored. 89 peach germplasms were selected to determine the trichomes lengths at fruit maturity. The analysis of the results suggested a weak correlation between trichome lengths and the expression levels of *Prupe.7G196500*. It was found that the trichome’s length reduced as peach fruit development progressed (Figure 6E). A combination of the results obtained suggested the transcript levels of *Prupe.7G196500* were inversely proportional to fruit growth (Figure 6D); therefore, we speculate that *Prupe.7G196500* may have a positive correlation with trichome development.

**Figure 6.**
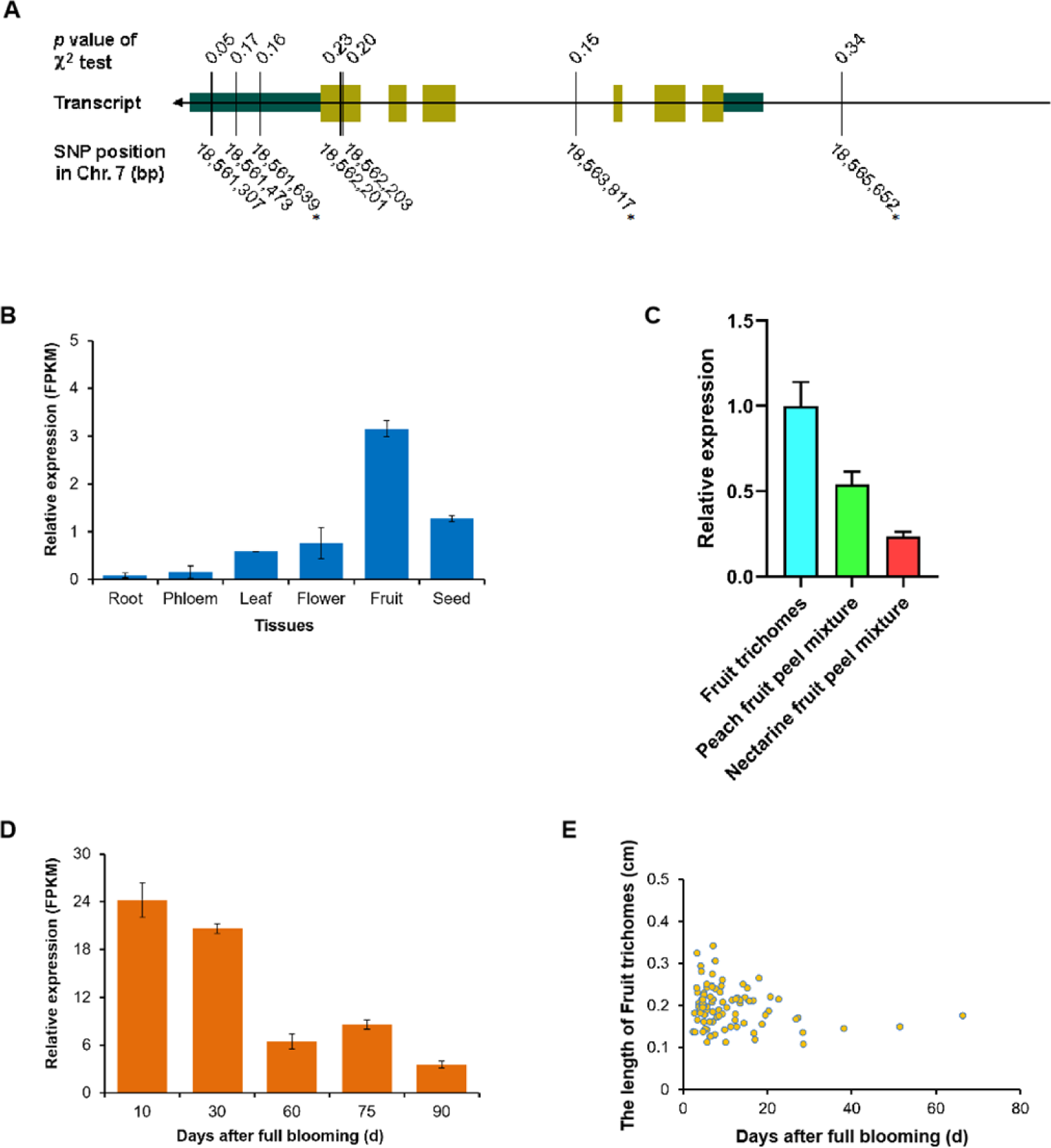
Prediction and analysis of the functions of *Prupe.7G196500* in peach. **(A)** Genotypic variation of *Prupe.7G196500* in nectarine and peach and the association of this variation with the phenotype. **(B)** Analysis of the tissue-specific expression of *Prupe.7G196500*. **(C)** Analysis of the tissue-specific expression of *Prupe.7G196500* in nectarine and peach fruits at the immature stage. **(D)** Analysis of the expression patterns of *Prupe.7G196500* at different time points during fruit development. **(E)** Analysis of the correlation between the length of fruit trichomes and the expression of *Prupe.7G196500* during the ripening of hairy peach fruits.

The *Prupe.7G196500pro::GUS* construct was introduced into *Arabidopsis* plants to evaluate the function of *Prupe.7G196500*. Specific GUS expression was detected in the trichomes of the true leaves of 10-day-old *Arabidopsis* seedlings under normal growth conditions (Figure 7A). Additionally, GUS expression was detected in the trichomes of the true leaves and epidermal cells post-treatment with 40 μM JA (Figure 7B), indicating that JA could induce the expression of *Prupe.7G196500*. Next, we validated the role of *Prupe.7G196500* in trichome formation by introducing the *p35S::Prupe.7G196500* overexpression construct into *Arabidopsis* seedlings. After screening two overexpression lines, *35S::Prupe.7G196500-3* and *-4,* were identified (Figure 8G). The number of trichomes in the true leaves of *35S::Prupe.7G196500-3* and *-4* seedlings significantly increased compared with that of the 10-day-old WT plants under normal growth conditions (Figure 8A, B, C, and H). Similarly, the number of trichomes in the true leaves of the *35S::Prupe.7G196500-3* and *-4* seedlings were significantly higher than in the WT plants upon 40 μM JA treatment (Figure 8D, E, F, and H).

**Figure 7.**
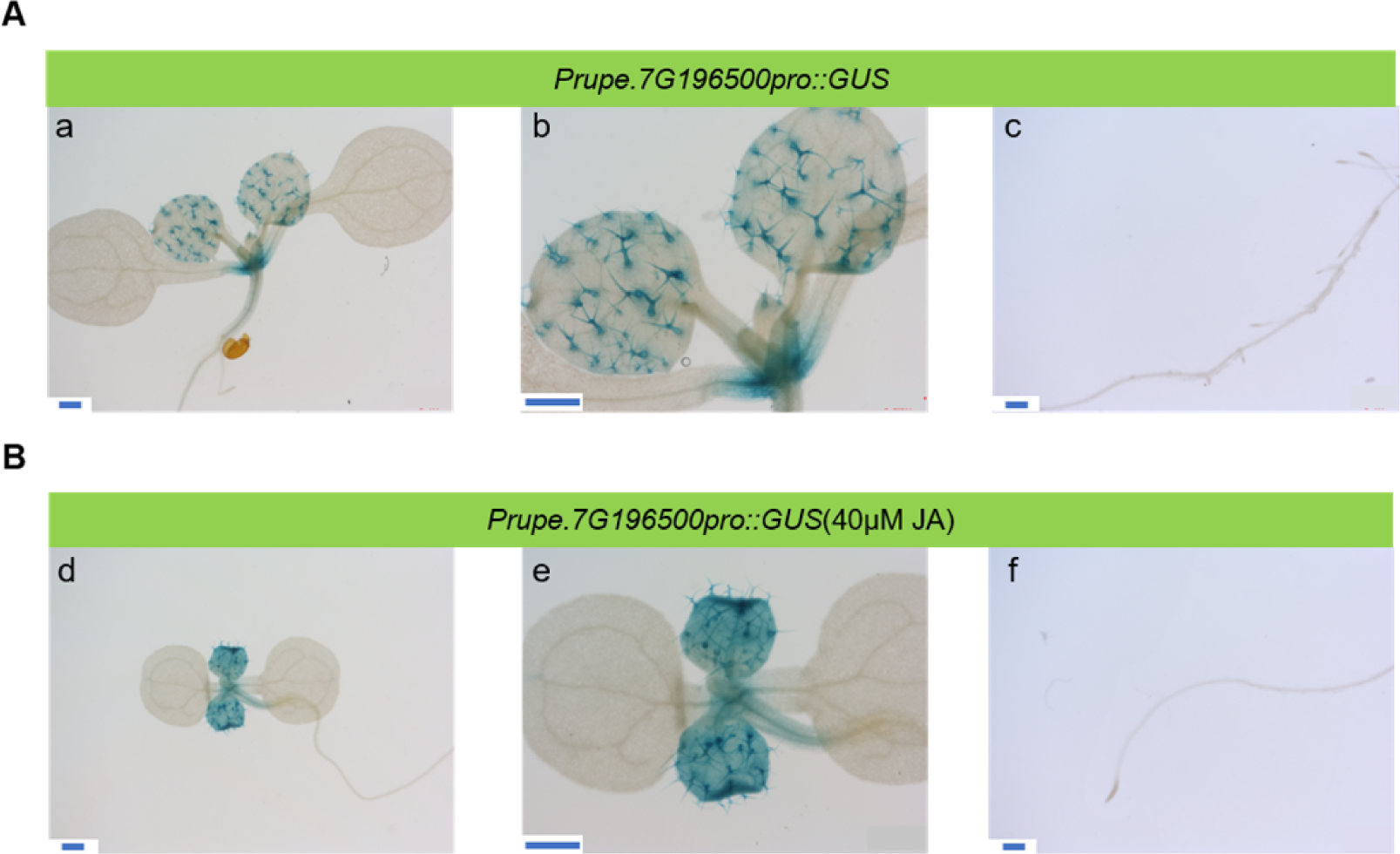
Analysis of the expression patterns of the marker genes representative of fruit trichomes in *Arabidopsis*. To detect the expression patterns of the marker genes representative of fruit trichomes, transgenic lines of *Arabidopsis* expressing the reporter gene GUS driven by the promoter of each gene were generated. **(A)** The GUS signals were detected in the true leaves of 10-day-old seedlings under mock conditions. Scale bar: 500 µm. **(B)** The levels of GUS were detected in the true leaves of 10-day-old seedlings after processed by 40 μM JA. Scale bar: 500 µm.

**Figure 8.**
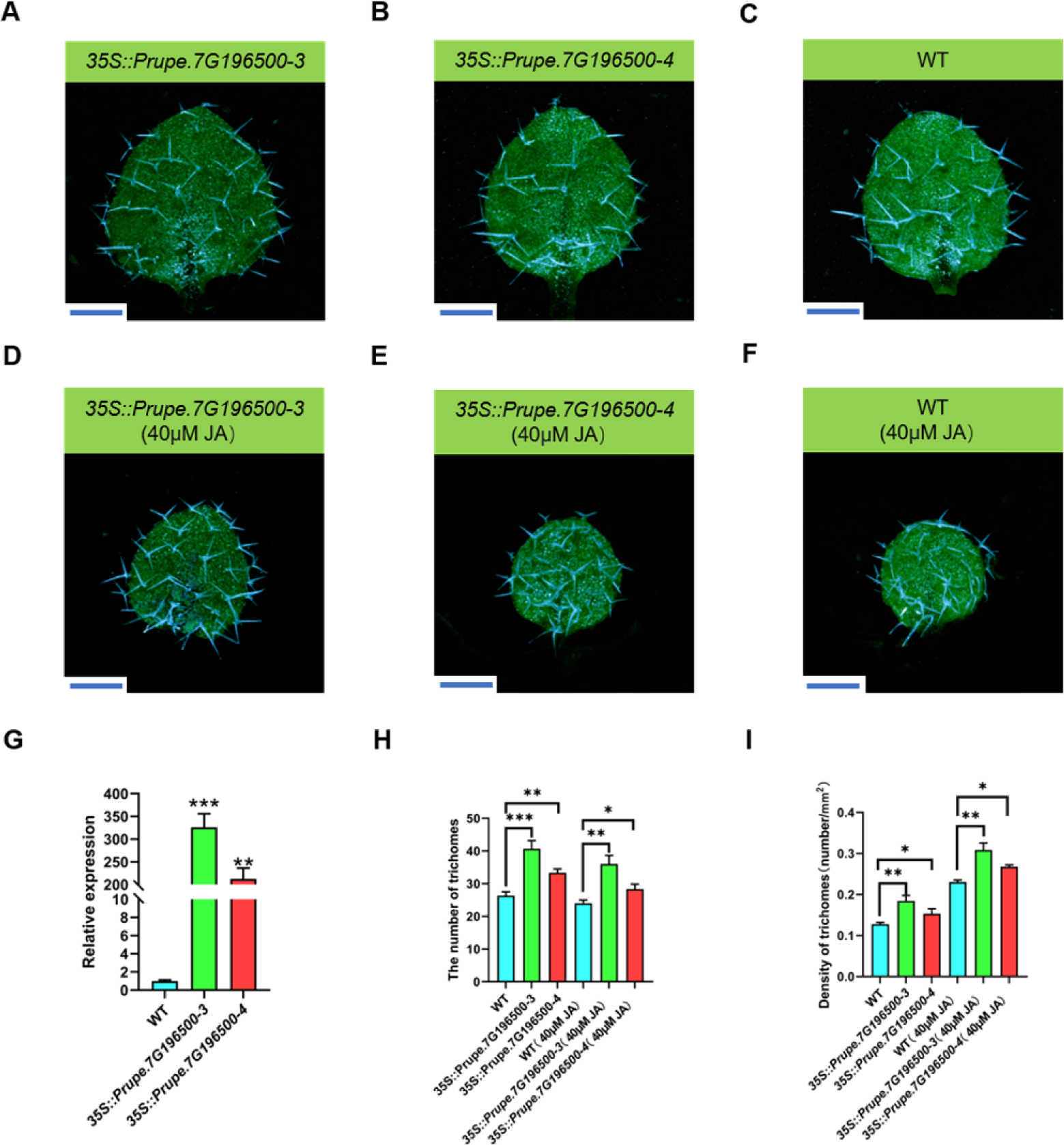
Phenotype analysis of trichomes growth in transgenic *Arabidopsis*. **(A – C)** Observation of the trichomes of the true leaves of 10-day-old transgenic and WT *Arabidopsis* plants under normal growth conditions. Scale bar: 500 µm. **(D – F)** Observation of the trichomes of the true leaves of 10-day-old transgenic and WT *Arabidopsis* plants after 40 μM JA treatment. Scale bar: 500 µm. **(G)** qPCR analysis of the relative expression levels of the representative marker gene, *Prupe.7G196500* in the *35S:: Prupe.7G196500*-3/4 overexpressing and WT *Arabidopsis* plants. * p < 0.05, ** p < 0.01, *** p < 0.001; one-way ANOVA Vs WT. **(H)** Analysis of the number of trichomes in the *35S::Prupe.7G196500*-3/4 overexpressing and WT *Arabidopsis* plants after treatment with or without 40 μM JA. * p < 0.05, ** p < 0.01, *** p < 0.001; one-way ANOVA Vs. WT. **(I)** Analysis of the density of trichomes in the *35S:: Prupe.7G196500*-3/4 overexpressing and WT *Arabidopsis* plants after treatment with or without 40 μM JA. * p < 0.05, ** p < 0.01, *** p < 0.001; one-way ANOVA Vs. WT.

Furthermore, the density of trichomes on *35S::Prupe.7G196500-3* and *-4* seedlings under normal conditions and post-treatment with 40 μM JA were statistically significant than that of the WT (Figure 8I). Notably, the *35S::Prupe.7G196500-3* and *-4* seedlings treated with 40 μM JA exhibited significantly higher trichome densities in the true leaves than the overexpression seedlings grown under normal conditions. Similarly, the trichome density of the true leaves of the WT seedlings post-treatment with 40 μM JA increased significantly than the WT seedlings grown under normal conditions, with a relative amplitude similar to that of the overexpression seedlings (Figure 8I). This indicated that the increase under normal growth conditions was not due to an enhancement in the JA signaling pathway but the overexpression of *Prupe.7G196500*. Therefore, it can be speculated that JA treatment induced the expression of *Prupe.7G196500* and trichome development, which may be independent of JA signaling.

### *Prupe.7G196500* plays a role in providing resistance to drought-induced stress in Arabidopsis

The GO enrichment analysis of DEGs from nectarine and peach showed that nectarine was more susceptible to stress but more resistant to drought than peach. A DEG, *Prupe.7G196500,* was identified in fruit trichomes (cluster 10), with enriched functions indicated as “response to wounding” and “response to jasmonic acid” (Supplemental Table S2). The role of *Prupe.7G196500* in mediating abiotic stress was verified by subjecting *Prupe.7G196500pro::GUS* expressing seedlings to various stress treatments. After treatment with 100 μM mannitol, the GUS signal was significantly higher than that in the control group, suggesting that drought stress could induce the expression of *Prupe.7G196500* (Supplemental Figure S8). Then, when the WT and overexpressing seedlings were subjected to drought-induced stress on MS medium, we found that the growth-related phenotypes of both types of seedlings cultured under standard conditions were almost similar (Supplemental Figure S9A and B), but under drought stress conditions, the leaves of the overexpression lines were larger and greener and developed a longer taproot than in the WT (Supplemental Figure S9C and D). The seedlings grown on MS medium were then transplanted into nutrient-rich soil for further observation. Consistent phenotypic growth of WT and overexpression seedlings under normal growth conditions were observed (Supplemental Figure S10A, C, E, G). Interestingly, no significant differences detected in the growth-associated phenotypes of the WT and overexpression seedlings after 14 days to drought exposure (Supplemental Figure S10B and D). However, the WT seedlings failed to survive after 21 days of dehydration, while the overexpression seedlings survived (Supplemental Figure S10F and H). Therefore, it can be proposed that *Prupe.7G196500* may be involved in regulating the tolerance of plants to drought-induced stress.

## DISCUSSION

In this study, the transcriptome information of peach and nectarine fruits at seven DAF was compared and analyzed using ST sequencing technology. Different cell clusters were classified and defined based on the tissue-type of the cell populations, the spatial expression patterns of marker genes, and the known functions of the homologs of these genes in other species (Figure 1), which enriched the available resources regarding the annotation of the cell type. In contrast with the cell populations of nectarine fruits, a unique cell population was identified in peach fruits and annotated as fruit trichomes (cluster 10) (Figure 1B). Then, a marker gene each for the exocarp and an endocarp was selected for ISH to demonstrate the reliability of the cell cluster definitions made (Figures 2 and SupplementalS3). The marker genes for fruit trichomes (cluster 10) were characterized as functionally influential genes potentially involved in forming trichomes in peach fruits (Figure 3). Notably, *Prupe.7G196500* was identified to be crucial for regulating trichome development. *Prupe.7G196500pro::GUS* was explicitly expressed in the trichomes of the true leaves of *Arabidopsis* (Figure 7). The number and density of the trichomes of the true leaves in the *35S::Prupe.7G196500* overexpression seedlings were significantly higher than those in the WT (Figure 8). Further analysis showed that the expression of *Prupe.7G196500* could be enhanced by drought-induced stress, suggesting that it may play a role in peache response to drought-induced stress (Supplemental Figures S8 – 10).

### Nectarine and peach fruits respond to stress in varied ways

Trichomes are highly differentiated epidermal cells that play essential developmental roles and act as the first line of defense against abiotic and biotic stresses (Hauser, 2014). Trichomes directly protect plants against sunlight, heat, and UV radiation and indirectly affect plant transpiration, water use, and photosynthetic efficiencies (Bickford, 2016). Available reports demonstrated their role in conferring mechanical barriers to herbivores (Furstenberg-Hagg et al., 2013). The most apparent phenotypic difference between peach and nectarine fruits is the presence or absence of trichomes (Vendramin *et al*., 2014), suggesting that they may also vary in their responses to environmental stress. The GO enrichment analysis of the DEGs identified in nectarine and peach showed that the up-regulated DEGs enriched in nectarine were related to defense response, mainly including “response to biotic stimulus,” “defense response,” and “saponin biosynthetic process” (Figures 4 and S3), which proves that nectarine was more susceptible to stress than peach. In contrast, the down-regulated DEGs in nectarine fruits were mainly related to “response to water deprivation,” “abscisic acid-activated signaling pathway,” and “response to abscisic acid” (Figures 4 and S3), indicating that nectarine exhibited a better water retention capacity which may be due to the dense and waxy layer covering the exocarp, which also makes the nectarine fruits appear glossy and smooth (Yang *et al*., 2022). Consistent with the GO enrichment analysis results, the TF-regulatory network of DEGs in nectarine and peach fruits also proved that they possessed different developmental processes. Altogether, nine key TFs (including three WRKY, three AP2/ERF-ERF, two bZIP, and one HB-HD-ZIP TF) were identified among the DEGs, (Figure 5). These TFs are essential regulators of biotic and abiotic stresses in plants (Ambawat *et al*., 2013; Droge-Laser *et al*., 2018; Erpen *et al*., 2018; Wani *et al*., 2021; Xie *et al*., 2019). These observations suggested that nectarine and peach fruits showed significant differences in their reactions to stress during the early period of development.

### *Prupe.7G196500* positively correlate with trichomes development in peach fruits

This study used several peach and nectarine germplasms resources for genotypic identification. Seven loci were identified as mutations in the mRNA and promoter of *Prupe.7G196500*, among which the SNP located at the 3’-UTR (Chr7:18,561,307 bp) most correlated with trichome development in peach fruits (Figure 6A). *Prupe.7G196500* was expressed in various tissues of peach plants and at different developmental stages of the fruits. The results showed that *Prupe.7G196500* was mainly expressed in the trichomes of peach fruit during the early developmental stage (Figure 6B – D). Further, to explore a possible correlation between the length of trichomes and the expression levels of *Prupe.7G196500* during the ripening process of hairy peach fruits, the lengths of the trichomes on fruit maturity were measured and analyzed, which suggested that the correlation was weaker but the trichomes shortened as the peach fruits grew and developed (Figure 6E). More significantly, *Prupe.7G196500*, which was highly expressed in the peach fruit at the immature stage, positively regulated the development of trichomes and, subsequently, stress resistance through the presence of longer trichomes. However, gradual ageing of the exocarp in peaches decreased the sensitivity to drought-induced stress, reducing trichomes role in development and death. Therefore, as the fruit matured, the trichome development was inhibited by repressing *Prupe.7G196500* (Figures 6D and E). Based on the above analysis, we proposed that *Prupe.7G196500* may positively correlate with trichome development.

### *Prupe.7G196500* positively regulates *Arabidopsis* trichome development independent of the JA signaling pathway

The peach trichomes are unbranched, and each develops from a single epidermal cell of the fruit, while those in *Arabidopsis* trichomes are usually branched and develop from a single cell at the leaf base (Hülskamp et al., 1994). Therefore, although both are non-glandular and unicellular, they possess distinctive features. *Arabidopsis* is a model plant for studying the development of non-glandular trichomes; significant progress has been made in the molecular genetic basis of the pattern formation in non-glandular trichomes, especially by JA, a model based on the WD-Repeat/bHLH/MYB complexes has been proposed (Pesch and Hulskamp, 2009; Qi *et al*., 2011). In contrast, the regulation of the development of non-glandular trichomes in peaches remains poorly understood. The pubescence of peach fruits was found to be controlled by a single “G” locus, which was mapped to LG 5, and the nectarine trait was recessive to the peach trait (Dirlewanger et al., 1998). The MYB TF *PpMYB25* (*ppa023143m*) has been identified as a candidate for the “G” site, and the insertion of an LTR retrotransposon in its third exon resulted in the glabrous phenotype (Vendramin *et al*., 2014). Another MYB TF, *PpMYB26*, located downstream of *PpMYB25*, also played a crucial role in trichome formation in peach fruits (Yang *et al*., 2022). The glabrous phenotype of nectarine fruits may be due to the insertion of a retrotransposon in the third exon of *PpMYB25*, repressing the downstream *PpMYB26* and other regulatory genes. Although some progress has been made in understanding the mechanism of the formation and development of trichomes in peaches, the underlying regulatory network remains largely unclear. This study identified *Prupe.7G196500* gene, that is responsive to JA-mediated regulation. The *Prupe.7G196500pro::GUS* construct was detected to be specifically expressed in the trichomes of the true leaves of *Arabidopsis* (Figure 7A) and was induced by JA (Figure 7B). Overexpression of *Prupe.7G196500* in *Arabidopsis* enhanced the density of trichomes on the true leaves under both normal growth conditions and post-JA treatment (Figure 8I). Notably, the *35S::Prupe.7G196500-3/4* overexpression seedlings treated with JA demonstrated a significantly higher density of trichomes on the true leaves than the seedlings grown under normal conditions. Similarly, compared with WT seedlings grown under normal conditions, the density of the trichomes in true leaves of WT seedlings treated with JA also increased significantly, with a relative amplitude similar to that of the overexpression seedlings (Figure 8I). Hence, it can be speculated that the increase in the trichome density of overexpression seedlings under normal growth conditions may not be caused by an enhancement in the JA-based signaling pathway, but by the overexpression of *Prupe.7G196500*.

### *Prupe.7G196500* enhances drought-induced stress tolerance in plants

In this study, the homologs of *Prupe.7G196500* in *Arabidopsis* were found to be *AtSSL4* and *AtSSL5* with high homology. Furthermore, the phylogenetic classification showed it was closely related to *AtSSL5* (Supplemental Figure S6). *AtSSL4* and *5* belong to the defense-induction-associated genes in plants and encode the SSL proteins with a magnified expression in plants subjected to external stress (Sohani *et al*., 2009). The GUS signal in the *Prupe.7G196500pro::GUS* expression seedlings was significantly enhanced after drought exposure (Supplemental Figure S7). The growth-related phenotypes of the WT and *35S::Prupe.7G196500* overexpressing seedlings cultured under average growth and drought on MS medium were observed to be almost similar (Supplemental Figure S8A and B). However, under drought-induced stress, the leaves of overexpressing seedlings were larger and greener, and the taproots were longer than those of the WT (Supplemental Figure S8C and D). The *Arabidopsis* seedlings were later moved from the MS medium to a nutrient-rich soil for further observations. The growth of the WT and overexpressing phenotypes were consistent under normal growth conditions (Supplemental Figure S9A, C, E, G). Interestingly, no significant difference in growth was observed in the WT and overexpressing seedlings after 14 days of exposure to drought (Supplemental Figure S9B and D). However, the WT seedlings could not survive after 21 days of drought exposure, but the overexpressing seedlings survived (Supplemental Figure S9F and H). Therefore, it can be postulated that *Prupe.7G196500* has a similar function to *AtSSL4* and *5*, that enhances drought-induced stress tolerance in plants.

In summary, owing to the lack of information regarding the identification of gene expression patterns in specific tissues of peach fruits at a high resolution, culminated with the lack of the resources reported in previous studies, ST sequencing was used to map the spatial expression of genes in the peach and nectarine fruits at the early stage of development. Additionally, the marker gene resources available for tissue/cell types annotation and analysis were enriched for subsequent reference. A comparative study of the transcriptomic information of peach and nectarine fruits showed significant differences in response to stress in the early stages of development (Figures 4 and Supplemental S3). Notably, a novel gene, *Prupe.7G196500,* with high expression in the trichomes of peach fruits at the immature stage, was identified to positively regulate trichome development and enhance tolerance to drought-induced stress (Figures 6 – 8 and Supplemental S7 – 9). In conclusion, this study constructed the spatial expression maps of genes in the cells/tissues of peach and nectarine fruits and characterized specific potential marker genes for trichomes in peaches, laying the foundation for the further analysis of the regulatory network of trichome formation and development in peach.

## MATERIALS AND METHODS

### Plant materials and growth conditions

The *Arabidopsis* accession Columbia-0 (Col-0) was used as the wild type (WT). The seeds were surface sterilized with 5% NaOCl and germinated on vertical, half-strength Murashige and Skoog (1/2 MS) plates. All transgenic and WT plants were grown in a climate-controlled chamber at 22 ℃ and illumination at an intensity of 100 µmol photons m^−2^ s^−1^ under a 14 h light/10 h dark regime.

## Spatial transcriptomics

### Tissue sectioning and H&E staining

The samples of Nectarine_1, Nectarine_2, Peach_1, and Peach_2 fruits were directly embedded into the optimum cutting temperature (OCT) compound (Sakura Finetek, Torrance, CA, USA) in 1.5 mL centrifuge tubes, incubated in a CM 1950 cryostat slicer (Leica, Wetzlar, Germany) at −20 ℃ for 20 min, and sliced to a thickness of 10 μm. RNA was extracted from the peach slices, and the RNA integrity number (RIN) was assessed. Qualified samples with RIN values > 7 were stained with H&E (Sigma, MO, USA), incubated at 37 ℃ for 5 min, and scanned for imaging.

### Permeabilization and tissue optimization

Tissue samples prepared for ST were first permeabilized under optimum conditions using the Visium tissue optimization slides (Chromium, USA). The sections were placed into the capture areas of a Visium cassette (Chromium, USA) with 70 μL of modified permease in each well (2% w/v cellulase R10, 0.4% w/v macerozyme R10, 1% w/v pectinase, 1% w/v hemicellulase, and 0.4% snailase) and incubated for 24 min. The Visium cassette was removed, and the sections were visualized.

### Reverse transcription and library construction

After permeabilization, the captured RNA was reverse-transcribed into cDNA, and 10 µL of it was used to prepare the library after adaptor ligation.

### Processing of sequencing data and quantification of gene expression

The raw reads in the FASTQ format generated via high-throughput sequencing were processed with the Space Ranger software, version 1.2.0 (10× Genomics, USA) (Satija et al., 2015), and the sequences obtained were aligned with the cotton genome as a reference using the STAR software, version 2.7.10b (Dobin et al., 2013). The brightfield images of the sections were captured. The spatial barcode information was then used to align the reads to specific spots on the tissue sections using the images obtained by H&E staining as a basis. The total number of spots, the number of reads per spot, the number of genes detected, and the number of UMIs were determined to evaluate the quality of the sequencing reads. Finally, the gene-spot matrix was generated for gene expression analysis.

### Quality analysis and further processing of data

The Seurat package version 3.2.0 (Stuart et al., 2019) was used to perform the quality control step and process the data obtained using Space Ranger. Then, the SCTransform package (Hafemeister and Satija, 2019) was used to normalize and stabilize the variance in the ST data using a regularized negative binomial regression, and the data were stored in the SCT for further analysis.

### Dimensionality reduction and clustering

The batch effects on the ST data were corrected with the “batchelor” package in R software (Haghverdi et al., 2018), following the mutual nearest neighbors (MNN) approach proposed by Haghverdi *et al*. Then, the “FindClusters” function of the Seurat package was used to scale the gene expression levels and cluster the cells based on the MNN approach. This strategy ultimately removed the batch effects on the ST data and enabled the detection of subpopulations of cells. Finally, the “RunTSNE” function of the Seurat package, which is based on a two-dimensional t-distributed stochastic neighbor embedding (t-SNE) algorithm, was used to visualize the cells (van der Maaten, 2014).

### Differential gene expression and enrichment analysis

The “FindMarkers” function (test. use = MAST) of the Seurat package (Butler et al., 2018) was used to identify the DEGs; those with a |log_2_ fold change| > 0.58 and a P value < 0.05 were identified as the DEGs. Then, GO (Carbon et al., 2019) and KEGG (Kanehisa et al., 2008) pathway enrichment analyses of the DEGs were performed using the R software based on the hypergeometric distribution.

## RNA ISH

ISH of the mRNA was performed using the kit (Boster Biological Technology, Wuhan, China), following the instructions of the manufacturer. The peach and nectarine fruits were collected, embedded in paraffin, and cut into 5 μm thick sections. The sections were de-waxed using xylene and rehydrated using a series of alcohol gradients. Subsequently, 3% citric acid and concentrated pepsin (two drops) were added to the sections and incubated at 37 ℃ for 10 min to obtain the mRNA. The cells were incubated with a DIG-labelled RNA probe (BOSTER) overnight at 60 ℃ for mRNA hybridization, washed twice with PBS, blocked with serum, and labeled with alkaline phosphatase-conjugated anti-Digoxigenin antibody, anti-Dig-AP (BOSTER) at 37 ℃ for one h. The alkaline phosphatase activity of the cells was detected in the dark using a nitro-blue tetrazolium/5-bromo-4-chloro-3-inodyl-phosphate (NBT/BCIP) based chromogenic substrate solution (BOSTER). The sections were analyzed, and the images were captured using an Aperio VERSE 8 multifunctional tissue cell analyzer (Leica Biosystems, Wetzlar, Germany). The positive cells were stained bluish-violet. Primers or probes used in this assay are listed in Table S5.

### Construction of expression vectors

The expression vectors were constructed using the ClonExpress^®^ MultiS One-Step Cloning Kit (Vazyme Biotech, Nanjing, China). The sequence 2000 bp upstream of the start codon was PCR-amplified, and the purified PCR product was inserted into the pCAMBIA1305.1 vector to generate the construct for the promoter of *Prupe.7G196500*. To create the GFP-fusion expression vector for *Prupe.7G196500*, the full-length CDS of *Prupe.7G196500* (1077 bp) was PCR-amplified, and the purified PCR product was inserted into the pCAMBIA2300 vector. The sequences of the primers used are listed in Table S5.

### Transformation of *Arabidopsis*

The *Agrobacterium tumefaciens* strain GV3101 cells were transformed with the GFP-fusion expression and GUS reporter constructs via electroporation. These were then used to transform the *Arabidopsis* WT plants using the floral dip method (Zhang et al., 2006). For the selection of the transgenic plants, kanamycin was used to screen the pCAMBIA2300-GFP-*Prupe.7G196500* expressing plants, and hygromycin for the pCAMBIA1305.1-promoter-*Prupe.7G196500* expressing plants. Homozygous transgenic lines were used for all the experiments.

### GUS staining and histological analysis

Histochemical GUS staining was performed using the G3061 GUS staining kit (Solarbio^®^ Life Sciences, Beijing, China) according to the instructions provided and as previously described (Liu et al., 2022).

### GO enrichment analysis

GO enrichment analyses for the DEGs were conducted using the Metascape resource (http://metascape.org/) (Zhou et al., 2019).

### Accession numbers

The sequence data obtained in this study were uploaded to the Genome Database for Rosaceae (GDR; https://www.rosaceae.org) with accession number *Prupe.7G196500*. ST data are available at the following web addresses: (https://dataview.ncbi.nlm.nih.gov/?search=SUB12286374).

## ACKNOWLEDGMENTS

We are grateful to ABRC for the *Arabidopsis* seeds. This research was supported by the National Key Research and Development Program of China (No.2022YFD1200300).

## AUTHOR CONTRIBUTIONS

Conceptualization of the project: X.S. and K.C. Experimental design: X.S. Performance of some specific experiments: A.Q., K.C., Z.Z., Z.L., L.G., S.S., H.L., Y.Z., J.Y., Y.L., M.H., V.N., and Z.Z. Data analysis: A.Q., K.C., Z.Z., and X.S. Manuscript drafting: A.Q., Z.Z., Z.L., and S.X. Contribution to the editing and proofreading of the manuscript draft: V.N., and L.W. All authors have read and approved the final manuscript.

## DATA AVAILABILITY STATEMENT

All data supporting the findings of this study are available within the paper and within its supplementary data published online.

